# Uncovering novel functions of the enigmatic, abundant and active *Anaerolineae* in a salt marsh ecosystem

**DOI:** 10.1101/2024.08.27.609934

**Authors:** Paige E. Payne, Loren N. Knobbe, Patricia Chanton, Julian Zaugg, Behzad Mortazavi, Olivia U. Mason

**Affiliations:** Department of Earth, Ocean and Atmospheric Science, Florida State University, Tallahassee, FL, USA; Department of Biological Sciences, University of Alabama, Tuscaloosa, AL, USA; Australian Centre for Ecogenomics, School of Chemistry and Molecular Biosciences, University of Queensland, St Lucia, Queensland, AU; Dauphin Island Sea Lab, Dauphin Island, AL, USA

## Abstract

*Anaerolineae*, particularly uncultured representatives, are one of the most abundant microbial groups in coastal salt marshes, dominating the belowground rhizosphere, where over half of plant biomass production occurs. However, this class generally remains poorly understood, particularly in a salt marsh context. Here, novel *Anaerolineae* metagenome assembled genomes (MAGs) were generated from the salt marsh rhizosphere representing *Anaerolineales*, *Promineifilales,* JAAYZQ01, B4-G1, JAFGEY01, UCB3, and *Caldilineales* orders. Metagenome and metatranscriptome reads were mapped to annotated MAGs, revealing nearly all *Anaerolineae* encoded and transcribed genes required for oxidation of simple sugars to complex carbon compounds, fermentation, and carbon fixation. Further, the majority of *Anaerolineae* expressed genes involved in anaerobic and aerobic respiration and secondary metabolite production. The data revealed that the belowground salt marsh *Anaerolineae* in the rhizosphere are important players in carbon cycling, including degradation of simple carbon compounds to more recalcitrant plant material, such as cellulose, using a diversity of electron acceptors and represent an unexplored reservoir of novel secondary metabolites.

**Importance:** Given that coastal salt marshes are recognized as biogeochemical hotspots, it is fundamentally important to understand the functional role of the microbiome in this ecosystem. In particular, *Anaerolineae* are abundant members of the salt marsh rhizosphere and have been identified as core microbes, suggesting they play a particularly important functional role. Yet, little is known about the metabolic pathways encoded and expressed in this abundant salt marsh clade. Using an ‘omics based approach we determined that *Anaerolineae* are capable of oxidizing a range of carbon compounds, including simple sugars to complex carbon compounds, while also encoding fermentation and carbon fixation. Surprisingly, *Anaerolineae* encoded and transcribed genes involved in aerobic respiration, which was unexpected given the reduced nature of the salt marsh rhizosphere. Finally, the majority of *Anaerolineae* appear to be involved in secondary metabolite production, suggesting that this group represents an unexplored reservoir of novel and important secondary metabolites.

## Introduction

*Chloroflexota* represents a metabolically diverse clade with members found globally in a multitude of primarily anaerobic environments (1). Early cultivated representatives of *Chloroflexota* were shown to grow in multicellular filaments and utilize an obligately anaerobic fermentative metabolism (2). More recently, culture-independent methods, including metagenomic sequencing and assembly, have been utilized to study the *Chloroflexota* in diverse biomes, including anaerobic digesters (3,4), marine sponges (5), and deep sea sediments (6). These studies demonstrated that the *Chloroflexota* can utilize diverse carbon substrates (5) and can ferment (1,4,7,8). In addition, the metabolic capabilities of the *Chloroflexota* have been shown to be more diverse than previously thought, including the capacity for nitrite/nitrate reduction (8–10), sulfate/sulfite reduction (6,11), aerobic respiration (8–10,12), phototrophy (13), and carbon fixation (14,15).

An accumulation of microbial data from a diversity of salt marshes, including those in MA and AL in the United States and in the Netherlands, have established *Chloroflexota* as one of the most abundant microbes in the belowground salt marsh environment (16–21), regardless of sampling season and/or depth. In particular the *Anaerolineae* class is one of the most dominant groups in the salt marsh rhizosphere, and in some cases, has been identified as a core rhizosphere community member (18,20,21). Yet, the functional role these abundant microbes play in a salt marsh is unknown, making it challenging to interpret the long-standing and rich plant and chemical datasets that exist for these environments (e.g. 22,23). Given that coastal salt marshes can overlie millennial-aged deposits of organic matter (24–26), organic carbon transformations mediated by microbes, such as the abundant *Anaerolineae*, would influence the carbon stored (or consumed) in salt marsh sediments (27).

Further, members of phylum *Chloroflexota* have previously been demonstrated to encode biosynthetic gene clusters (BGCs), groups of genes located in close proximity that represent a pathway for biosynthesis of a secondary metabolite (28), such as non-ribosomal peptide synthases (NRPS), polyketide synthases (PKS), NRPS-PKS hybrids, and lanthipeptides (29). These secondary metabolites can have potential medical applications, including as antibiotics (30) and immunosuppressants (31). The production of secondary metabolites by microbes in salt marshes is largely uncharacterized, with a few cultivation studies suggesting this is an important process in these environments (32,33). The high diversity of poorly described, uncultured microbial taxa, but particularly the abundant *Anaerolineae*, within salt marsh soils makes this environment ideal for the discovery of novel secondary metabolites by mining metagenomic and metatranscriptomic data (16,34).

In our previous study, we hypothesized that the rhizosphere of *Juncus roemerianus* (henceforth referred to as *Juncus*) and *Sporobolus alterniflorus* (previously *Spartina alterniflora*, henceforth referred to as *Sporobolus*) collocated at the same elevation in a Gulf of Mexico salt marsh would host distinct rhizosphere microbial communities (21). Yet, we found that microbial communities were highly similar, with comparable species richness, with differences observed only in species abundances, and most notably in that of the abundant *Anaerolineae.* Whether oscillations in the abundance of *Anaerolineae* members is important when interpreting both plant and chemical data for this site, or in other global salt marshes, remained unclear given the lack of cultured representatives, or ‘omics studies that have led to new genomic information. To address this knowledge gap and determine the functional role of the abundant, yet uncultured *Anaerolineae* in a salt marsh we carried out metagenomic and metatranscriptomic sequencing on rhizosphere samples from both plant types and from two different depths. Metagenomic data was assembled and annotated, allowing us to extend beyond taxonomic information and evaluate the function of salt marsh *Anaerolineae* (21). Samples from each plant type at two depths were also selected for metatranscriptomic sequencing. To evaluate gene and transcript abundance metagenomic and metatranscriptomic reads were mapped to metagenome assembled genomes (MAGs).

## Materials and Methods

### Study Site

The Dauphin Island (30°15.43’N, 88°07.438’W) salt marsh sampled is dominated by *Sporobolus* and interspersed with *Juncus*. As reported in Mason et al. (2021), at this site, tides are diurnal with mean tidal amplitude of less than 0.5 m, with flooding occurring on every high tide (21). Samples were collected from the *Juncus* and *Sporobolus* rhizosphere at 0–2, 5–7 and 0–10 cm in May 2015, 2016, and 2017 and immediately frozen at -80 °C.

### DNA Extraction and Purification

As described in Mason et al (2021), DNA (and RNA) was co-extracted from 1 g of sediment, from which any remaining root material was removed (21). Specifically, a modified CTAB extraction buffer ((10% CTAB (hexadecyltrimethylammonium bromide), 1M NaCl and 0.5M phosphate buffer, pH 8) with 0.1M ammonium aluminum sulfate, 25:24:1 phenol:chloroform:isoamyl alcohol) and bead beating using a FastPrep-24 (MP Biomedicals) was used to extract DNA and RNA (36). For DNA, eight samples from the *Juncus* and *Sporobolus* rhizosphere at 0–2, 5–7 and 0–10 cm were selected. The first and second DNA extractions were combined after the ethanol wash with 50 μl EB buffer to maximize DNA yields. The QIAGEN AllPrep DNA/RNA Kit (QIAGEN) was used to purify DNA and RNA following the manufacturer’s protocol (21). For metatranscriptomics, RNA was extracted from four samples of *Sporobolus* and *Juncus* at 0–2 and 5–7 cm as described above. Only RNA with an RNA integrity number (RIN) (16S/23S ribosomal RNA (rRNA) ratio determined with the Agilent TapeStation) ≥ 8 (on a scale of 1–10, with 1 being degraded and 10 being undegraded RNA) was selected for sequencing.

### Metagenomic and Metatranscriptomic Sequencing and DNA Sequence Assembly

Metagenomic sequencing was done by JGI following their metagenome SOP 1064. Sequencing was carried out on the eight samples described above, using the Illumina NovaSeq Regular 270 bp fragment mode. The BBTools software package, and specifically BBDUk and BBMap, were used to trim and screen paired-end Illumina reads (BBTools software package, http://bbtools.jgi.doe.gov). Reads were then read corrected using bfc (ver. r181) with bfc -1 -s 10g -k 21 -t 10 out.fastq.gz. Reads with no mate pair were removed. Low quality reads were subsequently removed using Trimmomatic (ver. 0.36, ILLUMINACLIP:TrueSeq) (37). MetaSPAdes (ver. 3.13) (38) was used for read assembly with default parameters. Contigs less than 500bp were removed using BBMap (ver. 38.41, https://sourceforge.net/projects/bbmap/). Quality controlled reads for each sample were mapped onto their respective assemblies using CoverM ‘make’ (ver. 0.2.0, B. Woodcroft, unpublished, https://github.com/wwood/CoverM) and low quality mappings were removed with CoverM ‘filter’ (minimum identity 95% and minimum aligned length of 50%). Using UniteM (ver. 0.0.15, D. Parks, unpublished, https://github.com/dparks1134/UniteM) scaffolds for each sample were binned by providing each sample’s contigs and BAM files as input with a minimum contig length of 1,500 bp. Maxbin (ver. 2.2.4) (39), MetaBAT (ver. 0.32.5) (40) and MetaBAT2 (ver. 2.12.1) (41) were used as binning methods (max40, max107, mb2, mb_verysensitive, mb_sensitive, mb_specific, mb_veryspecific and mb_superspecific). MAGs were dereplicated using dRep (ver. 2.2.3, sa = 0.95) (42). MAG completeness and contamination was evaluated using CheckM (ver. 1.0.12) (43). Subsequent analyses were inclusive of *Anaerolineae* MAGs that were medium-quality (≥50% completeness and <10% contamination) and high-quality (>90% completeness and <5% contamination) (Table 1) based on standards developed by the Genomics Standards Commission· (44). Using blastn, MAGs were compared to the 16S rRNA gene iTag data presented in Mason et al (2021) (21,45). For metatranscriptomics, JGI sequenced and processed cDNA reads using their metatranscriptome SOP 1066.1. Specifically, sequencing on the four samples described above was done using the Illumina NovaSeq platform in 2x151 mode. As with metagenomic sequence data, the BBTools software package was used to trim and screen reads. Further, reads containing known spike-ins and ribosomal RNA were removed. Before mapping cDNA reads a second round of ribosomal RNA was subtracted using riboPicker (ver. 0.4.3) with the default settings (46). Metagenomes used for assembly are available under NCBI bioproject accessions PRJNA539070–539077. MAGs are linked to the appropriate metagenome BioProject and are available in NCBI under SAMN41984578-79, SAMN41997743-66, SAMN41997787-99, SAMN41997938-54. Metagenomes used for read recruitment are available under NCBI bioproject accessions PRJNA539070–539075. Metatranscriptomes used for read recruitment are available under NCBI bioproject accessions PRJNA570129, 570130, 570132, and 535873.

**Table 1.** Taxonomy and assembly statistics for the 55 metagenome-assembled genomes (MAGs). MAGs were assigned to the high quality tier if they had >90% completeness and <5% contamination. MAGs were assigned to the medium quality tier if they had ≥50% completeness and <10% contamination.

| MAG Name | GTDB Classification* |  |  | Completeness (%) | Contamination (%) | Assembly Size (bp) | Number of Contigs | Assembly n50 | GC (%) | Number of Genes | Quality Tier |
| --- | --- | --- | --- | --- | --- | --- | --- | --- | --- | --- | --- |
|  | Order | Family | Genus |  |  |  |  |  |  |  |  |
| S5_7_050615r2r1_44 | Anaerolineales | Anaerolineaceae | Brevefilum | 65 | 10 | 3061922 | 1132 | 2792 | 0.45 | 3453 | Medium |
| J5_7_050615r2r3_11 | Anaerolineales | DRKV01 | JAGUYX01 | 92 | 8 | 5456903 | 744 | 11192 | 0.56 | 5210 | Medium |
| J5_7_160517rDrA_9 | Anaerolineales | E44-bin32 | E44-bin32 | 84 | 4 | 3303073 | 451 | 10459 | 0.60 | 3322 | Medium |
| S5_7_050615r2r1_21 | Anaerolineales | E44-bin32 | E44-bin32 | 80 | 6 | 2748020 | 665 | 4902 | 0.60 | 2955 | Medium |
| J0_2_160517rDrC_23 | Anaerolineales | EnvOPS12 | JAABUE01 | 79 | 7 | 4910627 | 1062 | 5659 | 0.53 | 5081 | Medium |
| J5_7_050615r2r3_15 | Anaerolineales | EnvOPS12 | Fen-1038 | 78 | 6 | 2984423 | 643 | 5972 | 0.61 | 3273 | Medium |
| S0_2_050615r3r5_73 | Anaerolineales | EnvOPS12 | UBA12087 | 56 | 2 | 1626702 | 272 | 7431 | 0.45 | 1671 | Medium |
| S5_7_050615r2r1_18 | Anaerolineales | RBG-16-64-43 | JAFGKK01 | 92 | 4 | 3166828 | 554 | 7541 | 0.62 | 3112 | High |
| S0_2_050615r3r5_74 | Anaerolineales | UBA11579 | UBA11579 | 64 | 6 | 2918571 | 777 | 4357 | 0.56 | 3153 | Medium |
| J0_2_160517rDrC_13 | Anaerolineales | UBA11858 | B10-G9 | 84 | 3 | 2537166 | 178 | 25006 | 0.56 | 2470 | Medium |
| S0_2_050615r3r5_38 | Anaerolineales | UBA11858 | B10-G9 | 80 | 4 | 2666833 | 339 | 10397 | 0.55 | 2666 | Medium |
| S0_2_050615r3r5_75 | Anaerolineales | UBA11858 | UBA11858 | 67 | 6 | 3420869 | 1010 | 4211 | 0.52 | 3781 | Medium |
| S5_7_050615r2r1_35 | Anaerolineales | UBA11858 | B10-G9 | 71 | 6 | 2382063 | 721 | 3799 | 0.55 | 2711 | Medium |
| S5_7_050615r2r1_74 | Anaerolineales | UBA11858 | UBA11858 | 68 | 8 | 5262343 | 1216 | 5242 | 0.47 | 5456 | Medium |
| J0_2_160517rDrC_8 | Anaerolineales | UBA4823 | PFL25 | 91 | 7 | 4529468 | 259 | 35346 | 0.49 | 4142 | Medium |
| J5_7_160517rDrA_4 | Anaerolineales | UBA4823 | PFL25 | 92 | 7 | 4441680 | 152 | 59425 | 0.49 | 3958 | Medium |
| J5_7_160517rDrA_8 | Anaerolineales | UBA4823 | DSOS01 | 87 | 3 | 3798058 | 337 | 22966 | 0.46 | 3583 | Medium |
| S_170502SBrC_35 | Anaerolineales | UBA4823 | CAIQV01 | 56 | 8 | 2402046 | 963 | 2494 | 0.42 | 2804 | Medium |
| S5_7_050615r2r1_1 | Anaerolineales | UBA4823 | PFL25 | 94 | 7 | 4603686 | 149 | 73311 | 0.49 | 4101 | Medium |
| S5_7_050615r2r1_61 | Anaerolineales | UBA4823 |  | 51 | 1 | 2350257 | 94 | 52461 | 0.49 | 2143 | Medium |
| J0_2_160517rDrC_47 | B4-G1 | B4-G1 | JAFGED01 | 81 | 10 | 4966031 | 1058 | 5832 | 0.59 | 4989 | Medium |
| S5_7_050615r2r1_67 | B4-G1 | B4-G1 | JAFGED01 | 70 | 9 | 4409391 | 607 | 10503 | 0.60 | 4136 | Medium |
| J5_7_160517rDrA_21 | B4-G1 | SLSP01 | DSWT01 | 73 | 4 | 3577821 | 998 | 4117 | 0.57 | 3616 | Medium |
| S5_7_050615r2r1_22 | B4-G1 | SLSP01 | DSWT01 | 82 | 1 | 4343584 | 980 | 5283 | 0.57 | 4042 | Medium |
| S0_2_050615r3r5_26 | Caldilineales | Caldilineaceae |  | 87 | 1 | 5570871 | 833 | 11921 | 0.58 | 5064 | Medium |
| J5_7_050615r2r3_21 | JAAYZQ01 | JAAYZQ01 |  | 73 | 8 | 4824464 | 1466 | 3609 | 0.62 | 5081 | Medium |
| J5_7_160517rDrA_32 | JAAYZQ01 | JAAYZQ01 |  | 55 | 2 | 3885091 | 1145 | 3759 | 0.62 | 4156 | Medium |
| S5_7_050615r2r1_46 | JAAYZQ01 | JAAYZQ01 |  | 72 | 7 | 4940851 | 1229 | 4805 | 0.62 | 5088 | Medium |
| S5_7_050615r2r1_50 | JAAYZQ01 | JAAYZQ01 | JAMJFU01 | 64 | 4 | 3571616 | 302 | 17099 | 0.63 | 3319 | Medium |
| S5_7_050615r2r1_68 | JAAYZQ01 | JAAYZQ01 | JAFGIA01 | 52 | 2 | 3548520 | 1050 | 3832 | 0.67 | 3642 | Medium |
| S0_2_050615r3r5_150 | JAAYZQ01 |  |  | 57 | 9 | 4452285 | 2458 | 1886 | 0.65 | 5507 | Medium |
| J0_2_160517rDrC_63 | JAFGEY01 | JAFGEY01 | JAFGIE01 | 59 | 9 | 5127177 | 1574 | 3860 | 0.64 | 5267 | Medium |
| J5_7_160517rDrA_13 | JAFGEY01 | JAFGEY01 | JAFGIE01 | 94 | 7 | 6262856 | 666 | 15834 | 0.64 | 5531 | Medium |
| S_170502SBrC_24 | JAFGEY01 | JAFGEY01 | JAFGIE01 | 78 | 9 | 4728695 | 1374 | 3984 | 0.64 | 4948 | Medium |
| S5_7_050615r2r1_52 | JAFGEY01 | JAFGEY01 | JAFGIE01 | 63 | 4 | 5138476 | 681 | 12622 | 0.65 | 4520 | Medium |
| J_170502JBrBrA_11 | Promineifilales | Promineifilaceae | JAACFF01 | 92 | 4 | 5232048 | 391 | 26559 | 0.52 | 4653 | High |
| J_170502JBrBrA_12 | Promineifilales | Promineifilaceae | JAACFF01 | 92 | 9 | 5343781 | 862 | 8857 | 0.57 | 4981 | Medium |
| J0_2_160517rDrC_16 | Promineifilales | Promineifilaceae | JAACFF01 | 81 | 3 | 4011945 | 203 | 43119 | 0.52 | 3528 | Medium |
| J5_7_050615r2r3_13 | Promineifilales | Promineifilaceae | JAACFF01 | 90 | 9 | 5603865 | 712 | 11414 | 0.57 | 5039 | Medium |
| J5_7_050615r2r3_16 | Promineifilales | Promineifilaceae |  | 71 | 4 | 3757630 | 980 | 4388 | 0.52 | 3788 | Medium |
| J5_7_050615r2r3_23 | Promineifilales | Promineifilaceae | JAACFF01 | 79 | 6 | 4557045 | 876 | 7154 | 0.53 | 4525 | Medium |
| J5_7_160517rDrA_23 | Promineifilales | Promineifilaceae | JAACFF01 | 77 | 3 | 4819677 | 907 | 6714 | 0.57 | 4534 | Medium |
| J5_7_160517rDrA_24 | Promineifilales | Promineifilaceae | JAACFF01 | 73 | 3 | 3126033 | 170 | 43029 | 0.51 | 2776 | Medium |
| S_170502SBrC_11 | Promineifilales | Promineifilaceae | JAACFF01 | 91 | 4 | 5113956 | 701 | 10932 | 0.56 | 4669 | High |
| S_170502SBrC_5 | Promineifilales | Promineifilaceae | JAACFF01 | 84 | 3 | 4409282 | 244 | 34589 | 0.53 | 3859 | Medium |
| S0_2_050615r3r5_25 | Promineifilales | Promineifilaceae | JAACFF01 | 78 | 3 | 3268746 | 168 | 37141 | 0.52 | 2849 | Medium |
| S0_2_050615r3r5_28 | Promineifilales | Promineifilaceae | JAACFF01 | 86 | 2 | 4710700 | 784 | 11491 | 0.57 | 4359 | Medium |
| S0_2_050615r3r5_39 | Promineifilales | Promineifilaceae | JAACFF01 | 95 | 6 | 5165683 | 707 | 10471 | 0.58 | 4669 | Medium |
| S0_2_160517rDrB_21 | Promineifilales | Promineifilaceae | JAACFF01 | 84 | 3 | 4586319 | 908 | 6258 | 0.57 | 4403 | Medium |
| S0_2_160517rDrB_7 | Promineifilales | Promineifilaceae | JAACFF01 | 95 | 4 | 5165867 | 306 | 34843 | 0.52 | 4524 | High |
| S5_7_050615r2r1_14 | Promineifilales | Promineifilaceae | JAACFF01 | 94 | 4 | 5627990 | 427 | 35849 | 0.53 | 4916 | High |
| S5_7_050615r2r1_16 | Promineifilales | Promineifilaceae | JAACFF01 | 89 | 2 | 4761143 | 785 | 9798 | 0.56 | 4442 | Medium |
| S5_7_050615r2r1_64 | Promineifilales | Promineifilaceae | UBA11865 | 52 | 3 | 3770228 | 1236 | 3384 | 0.60 | 3946 | Medium |
| S5_7_050615r2r1_33 | UCB3 | UCB3 |  | 70 | 4 | 2935820 | 327 | 13039 | 0.70 | 2374 | Medium |
| S0_2_050615r3r5_48 | UCB3 |  |  | 77 | 3 | 2425299 | 299 | 10196 | 0.70 | 2019 | Medium |

\*All genomes were classified as: Bacteria; Chloroflexota; Anaerolineae (not shown).

### Microbial Taxonomic Assignment, Read Recruitment, and Genome Similarity

Average nucleotide identity (ANI) was calculate using anvi’o (47) to run PyANI (48) with the ANIblastall parameter and default settings. Taxonomies were assigned to each MAG using the Genome Taxonomy Toolkit (GTDB-Tk, ver. 2.3.0; with reference to GTDB R08-RS214) (49) with up to 120 bacterial single-copy marker proteins. Functional annotations were assigned using DRAM (ver. 1.1.1) (50) with the KEGG (51), UniRef90 (52), PFAM (53), dbCAN (54), and MEROPS (55) databases. RRAP (ver. 1.3.2) (56), which automates read recruitment using Bowtie2 (57) and SAMtools (58), was used both to calculate MAG abundance and activity, by mapping metagenomic and metatranscriptomic reads to whole MAGs, and gene abundance and activity, by mapping metagenomic and metatranscriptomic reads to DRAM- annotated genes within each MAG. Metagenomic and metatranscriptomic read recruitment of whole MAGs and DRAM-annotated genes were normalized as reads per kilobase per million (RPKM) using RRAP. BGC genes were identified in MAGs using antiSMASH (ver. 7.0) (59). Because not all genes identified as part of BGCs were annotated by DRAM, Prodigal (ver. 2.6.3, https://github.com/hyattpd/Prodigal) was used to predict all genes in each MAG. BGC genes were then mapped to the MAGs with genes predicted by Prodigal using RRAP.

### Gene and Transcript Annotation

Genes related to organic carbon oxidation were divided into two groups based on carbohydrate active enzyme (CAZyme) category. Briefly, the term CAZyme represents carbohydrate-active enzymes including glycoside hydrolases, glycosyltransferases, polysaccharide lyases, carbohydrate esterases, and enzymes carrying out auxiliary activities (www.cazy.org) (54). Genes in the first group were CAZyme-encoding genes involved in degradation of complex carbon substrates and include glycoside hydrolases (EC: 3.2.1.- and 2.4.1.-) and a limited number of carbohydrate esterases (EC: 3.5.1.41). The second group contained genes related to monosaccharide oxidation as well as oxidation of polyphenolics, a secondary metabolite of plants (60). Genes related to carbon fixation were selected and grouped according to the KEGG modules for carbon fixation in prokaryotes. Genes utilized in fermentation were identified as those which acted upon pyruvate or an intermediary to eventually result in one of the major products of fermentation produced by *Anaerolineae* in culture: acetate, ethanol, lactate, formate, hydrogen, or succinate (61). Genes utilized in oxidative phosphorylation were limited to complex IV of the electron transport chain (ETC), which catalyzes the reduction of O_2_ to H_2_O during aerobic respiration (62). After mapping reads to annotated genes and transcripts, log1p-transformed RPKM values were summed and fold change was calculated to compare various groups to each other. In instances where one sample is being compared to multiple others (e.g. *Sporobolus* 5-7 cm against all plant species and depths), fold change was calculated between all possible pairs of samples and then averaged.

## Results

### Genome Summary, Taxonomy, and Relatedness of Salt Marsh Anaerolineae

In total, 55 *Anaerolineae* MAGs were chosen for analysis based on genome completion and contamination. Of these MAGs, 50 were considered medium-quality, with ≥50% completeness and <10% contamination, and five were considered high-quality, with >90% completeness and <5% contamination (Table 1). Taxonomic analysis revealed that the majority were classified as belonging to the *Anaerolineales* and *Promineifilales* orders (Table 1). Additional lineages included the orders JAAYZQ01, B4-G1, JAFGEY01, UCB3, and *Caldilineales* (Table 1). Of the 55 MAGs, 46 were assigned taxonomy to the genus level (Table 1). In six of these MAGs, including the dominant *Anaerolineales* and *Promineifilales*, 16S rRNA genes were identified and all were highly similar (bit score >100) to *Anaerolineae* 16S rRNA gene iTag data presented in Mason et al (2021) (21) (Figures 1 and 2). Two MAG 16S rRNA genes were most similar (bit score >370) to *Anaerolineales* that were identified as members of the core salt marsh rhizosphere microbiome (21).

**Figure 1.**
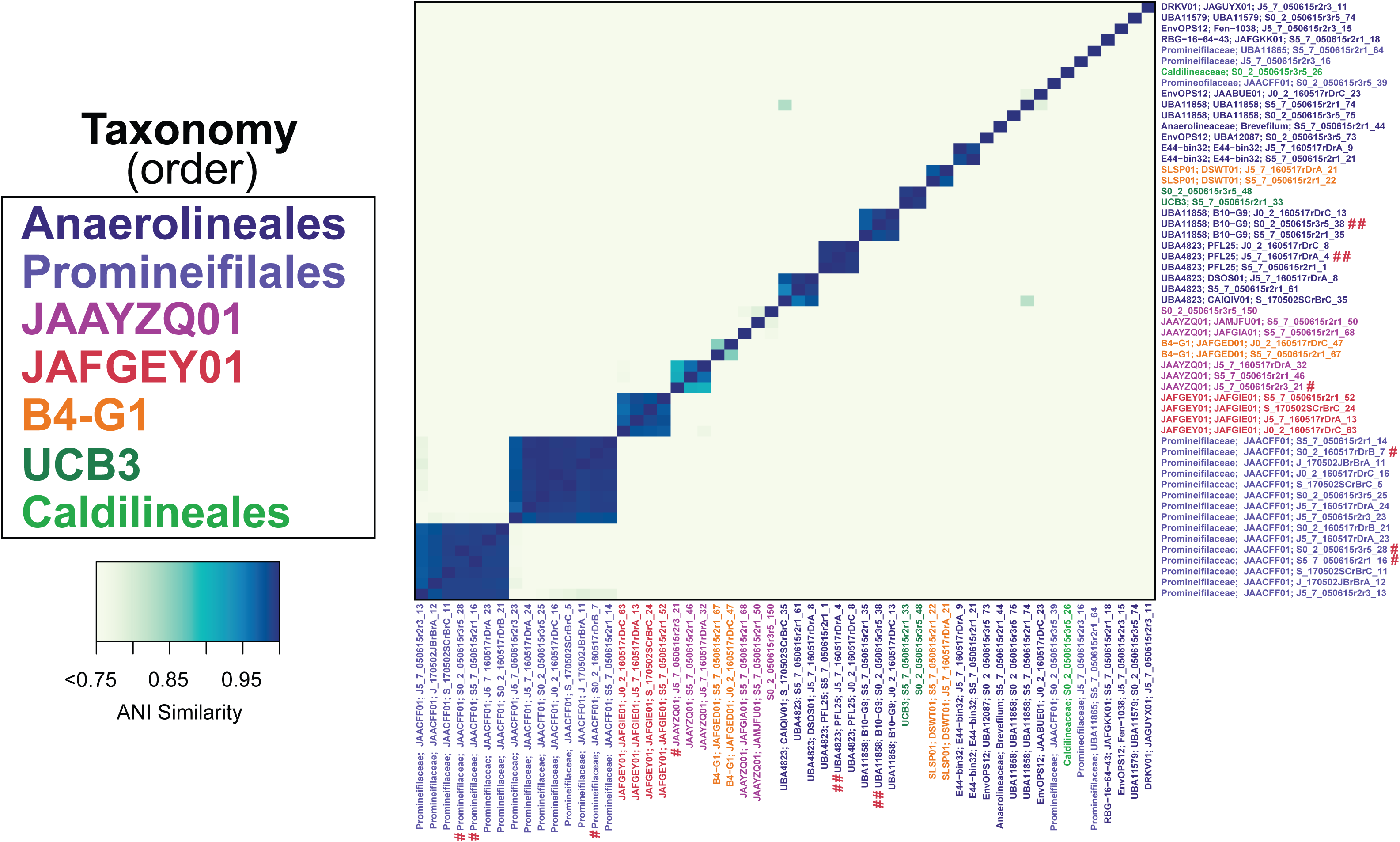
Average nucleotide identity (ANI) heatmap of the 55 *Anaerolineae* MAGs. Only ANI greater than 75% is shown. MAGs are color coded by taxonomic classification at the order level. In this figure # indicates a MAG 16S rRNA gene that is highly similar (bit score >100) to Anaerolineae 16S rRNA gene iTag data presented in Mason et al (2021) (21), while ## indicates MAG 16S rRNA genes were most similar (bit score >370) to core *Anaerolineales* in that same study.

**Figure 2.**
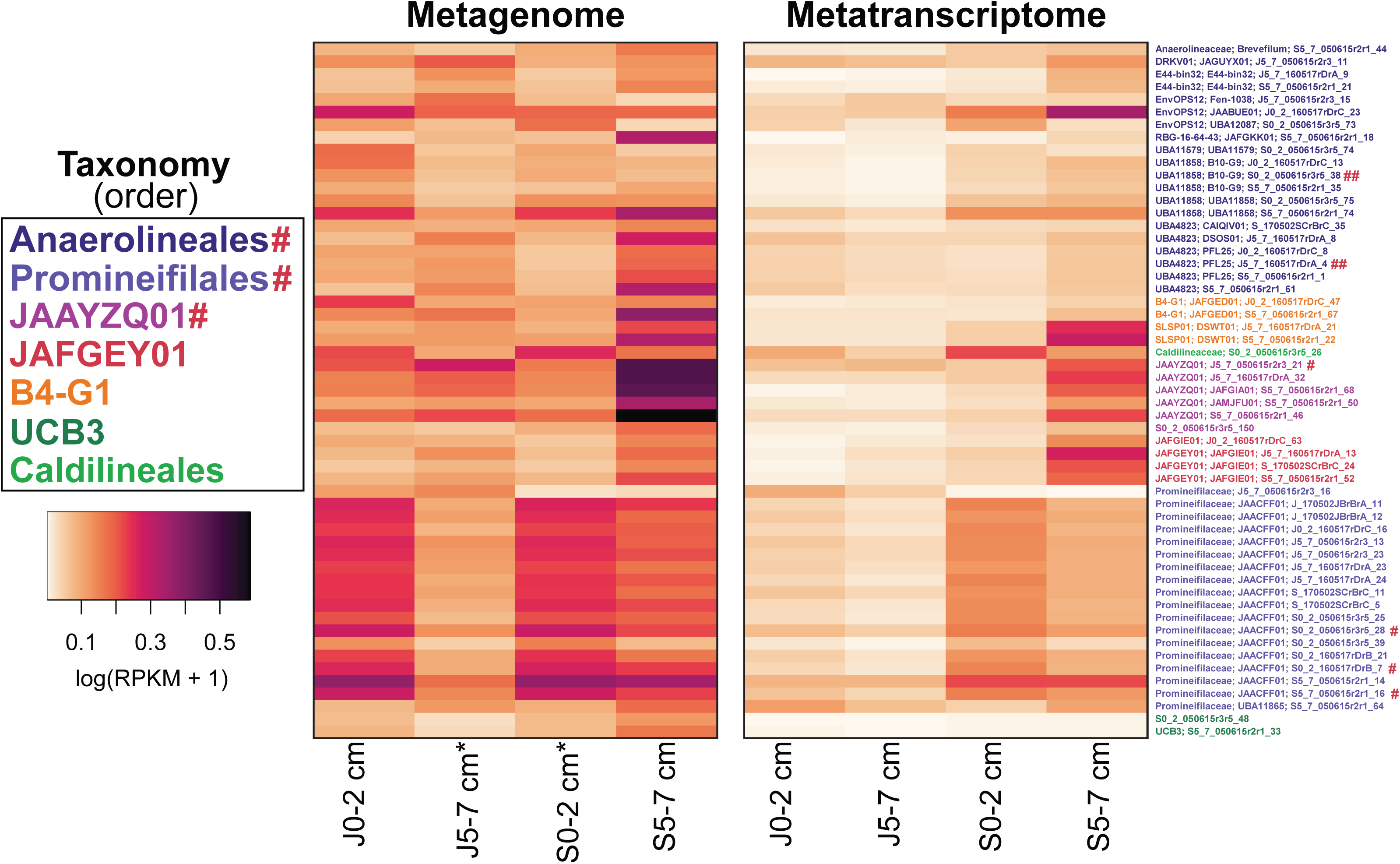
Microbial abundances determined by mapping metagenomes and metatranscriptomes from both plant species (*Juncus* and *Sporobolus*) and depths (0–2 cm and 5–7 cm) to the 55 *Anaerolineae* MAGs. Log1p-transformed RPKM values are shown. MAGs are color-coded by taxonomic classification at the order level. An asterisk (*) indicates that RPKM values from two metagenomes have been averaged together prior to transformation. In this figure # indicates a MAG 16S rRNA gene that is highly similar (bit score >100) to Anaerolineae 16S rRNA gene iTag data presented in Mason et al (2021) (21), while ## indicates MAG 16S rRNA genes were most similar (bit score >370) to core *Anaerolineales* in that same study.

ANI analysis showed that all MAGs were ≥59.1% in similarity (Figure 1). In addition, five MAGs classified as *Anaerolineales* and two groups in the *Promineifilales* order were observed to have ≥ 98.65% ANI, providing strong evidence that several *Anaerolineales* and *Promineifilales* within these groups likely belong to the same species (Figure 1) (63).

### Anaerolineae Abundance Based on Metagenomic and Metatranscriptomic Read Recruitment to Genomes

Microbial abundances were calculated by recruiting raw metagenomic and metatranscriptomics reads to MAGs. Metagenomic read mapping revealed that the five *Anaerolineae* MAGs with the highest read recruitment were classified as JAAYZQ01 and *Promineifilales* (Figure 2). In contrast, the five genomes that recruited the highest total number of RNA reads across all metatranscriptome samples included one MAG each from the orders *Anaerolineales*, JAAYZQ01, and *Caldilineales*, and two MAGs from the order *Promineifilales* (Figure 2). When RPKM values of all MAGs were summed together for both grasses and fold change was calculated, all MAGs recruited 0.43-fold more reads from *Sporobolus* metagenomes than from *Juncus* metagenomes, and 1.60-fold more reads from *Sporobolus* metatranscriptomes than from *Juncus* metatranscriptomes (Figure 2). Overall, the highest abundances (RPKM values) were MAGs from *Sporobolus* patches at 5–7 cm depth (Figure 2).

### Abundance and Expression of Carbon Utilization Genes by Substrate or Pathway

In general, the *Anaerolineae* MAGs analyzed here encoded genes allowing for utilization of a diverse range of organic carbon substrates, including both monosaccharides and more complex carbohydrates (Figure 3). For example, non-CAZyme-encoding genes involved in the consumption of carbon substrates including glucose, galactose, and xylose were abundant and highly expressed (Figure 3A). *Promineifilales* recruited the most total reads of non-CAZyme-encoding genes with total DNA RPKM of 1040.52 and a total RNA RPKM of 288.36, followed by the *Anaerolineales* (Figure 3A). When read recruitment from all MAGs across all samples was summed by substrate, glycolysis genes had the highest total read recruitment (RPKM values of 1552.80 DNA and 542.89 RNA). Complete glycolysis pathways were observed in 23/55 MAGs, which was inclusive of all observed orders except UCB3 (Figure 3C). An additional 24/55 MAGs were found to possess near-complete glycolysis pathways, or 7–8 steps out of 9, encompassing all observed orders (Figure 3C). *Promineifilales* had the highest total glycolysis gene read recruitment (RPKM of 662.26 DNA and 337.47 RNA) followed by the *Anaerolineales*. Genes encoding galactose degradation were also abundant, but less so than those involved in glycolysis when read recruitment from all MAGs was summed across all samples (Figure 3A). While metagenomic read recruitment of non-CAZyme-encoding genes was only 0.44-fold higher in *Sporobolus* patches than in *Juncus*, metatranscriptomic read recruitment was 2.03-fold higher in *Sporobolus* patches (Figure 3E). All MAGs recruited the greatest number of reads summed across all substrates from *Sporobolus* 5–7 cm samples, with total metagenomic read recruitment that was 0.68-fold higher compared to all other samples (Figure 3E). Similarly, total metatranscriptomic read recruitment to MAGs was 1.71-fold higher in *Sporobolus* 5–7 cm samples compared to the other plant type and depths (Figure 3E). The greatest DNA and RNA read recruitment from *Sporobolus* 5–7 cm was observed with every pathway that was analyzed; therefore, this observation is not repeated in the subsequent results.

**Figure 3.**
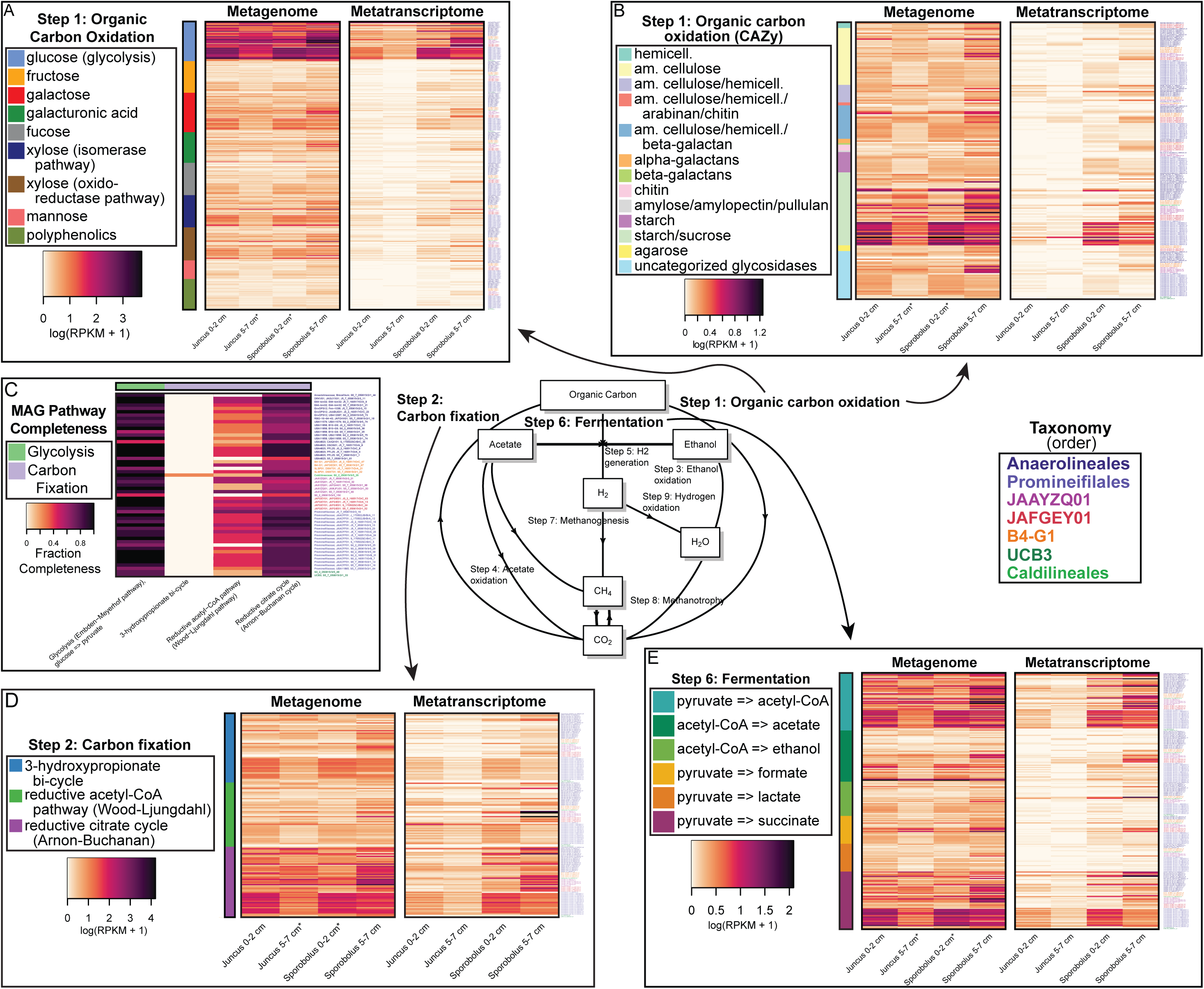
Gene and transcript abundances determined by mapping metagenome and metatranscriptome reads from both plant species (*Juncus* and *Sporobolus*) and depths (0–2 cm and 5–7 cm) to annotated genes and transcripts from the 55 *Anaerolineae* MAGs. Data shown is RPKM values that has been log1p-transformed. Data includes organic carbon oxidation by non-CAZyme-encoding genes (A) and CAZyme-encoding genes (B), carbon fixation (D), and fermentation (E) genes. Genes that can be considered part of more than one pathway have been included in all applicable pathways. Fraction completeness of certain pathways (C) was calculated by dividing the number of genes from a certain pathway encoded by each MAG (regardless of grass type or depth) by the total number of genes in the pathway. MAGs are color-coded by taxonomic classification at the order level. An asterisk (*) indicates that RPKM values from two metagenomes have been averaged together prior to transformation.

Although DNA and RNA read recruitment were highest for genes coding for degradation of sugars, there was strong evidence that *Anaerolineae* have the ability to degrade more recalcitrant forms of carbon. For example, across both plant species, all *Anaerolineae* MAGs encoded at least one CAZy gene (Figure 3B). Further, 50/55 MAGs transcribed at least one CAZyme degrading starch and sucrose (Figure 3B). Based on total recruited metagenomic and metatranscriptomic reads the greatest RPKM of CAZy genes was in the order *Promineifilales* (94.34 DNA and 22.33 RNA), followed by the *Anaerolineales* (Figure 3B). Recruitment of CAZy genes from metagenomes showed highest total recruited reads (RPKM of 102.33) for genes coding for enzymes used in degrading starch and sucrose, representing three genes involved in degrading both substrates, followed by amorphous cellulose degradation (Figure 3B). Metatranscriptomic read recruitment was similar, with genes involved in degrading starch and sucrose recruiting the highest number of reads (RPKM of 25.58), followed by cellulases (Figure 3B). These groups of CAZy genes included genes coding for cellobiose phosphorylase, sucrose phosphorylase, alpha-glucosidase, and beta-fructofuranosidase.

In addition to encoding and expressing genes used to degrade both simple and complex carbon sources, salt marsh *Anaerolineae* MAGs coded for carbon fixation. Total metagenomic read recruitment showed that *Promineifilales* recruited the most DNA reads from carbon fixation genes and pathways (RPKM of 486.23), followed by the *Anaerolineales* (Figure 3D). In contrast, carbon fixation genes and pathways encoded by JAAYZQ01 MAGs recruited the greatest number of metatranscriptomic reads (RPKM of 219.77), followed by *Promineifilales* (Figure 3D). Genes necessary for several alternative pathways to the Calvin Cycle were identified and transcribed, while no ribulose bisphosphate carboxylase/oxygenase (RuBisCO) genes were identified in the *Anaerolineae* genomes. The reductive citrate cycle was the most complete pathway in each MAG (Figure 3C). Specifically, 39/55 MAGs possessed ≥ 70% of the genes required to carry out the reductive citrate cycle. Pathway completeness was particularly high among members of the *Caldilineales*, *Anaerolineales*, and *Promineifilales* orders (Figure 3C). Of the three cycles highlighted in Figure 3D, the reductive citrate (Arnon-Buchanan) cycle recruited the greatest number of DNA reads (RPKM of 760.18) followed by the 3-hydroxypropionate bi-cycle (Figure 3D). Importantly, the majority (54/55) of MAGs lacked key genes required to fully carry out the 3-hydroxypropionate bi-cycle, and only one MAG belonging to the *Caldilineales* was found to encode and transcribe one of the four key genes (Figure 3C). The reductive citrate cycle also recruited the greatest number of RNA reads (RPKM of 346.68), followed by the reductive acetyl-CoA (Wood-Ljungdahl) cycle (Figure 3D).

Lastly, salt marsh *Anaerolineae* MAGs coded for anaerobic carbon consumption through fermentation. The MAGs of the order *Promineifilales* recruited the highest total number of DNA and RNA reads from fermentation pathways (324.26 DNA and 97.17 RNA), followed by the *Anaerolineales* (Figure 3E). The capacity of the *Anaerolineae* to carry out each of the fermentative pathways shown in Figure 3E was not uniform. The fermentative pathway pyruvate to acetyl-CoA, a precursor to both acetate and ethanol formation, recruited the greatest number of metagenomic and metatranscriptomic reads (249.87 DNA and 115.49 RNA). Similarly, the greatest number of MAGs (54/55) encoded and transcribed genes used to convert pyruvate to acetyl-CoA, including at least one representative from each order. This included genes for direct conversion of pyruvate to acetyl-CoA as well as indirect conversion of pyruvate to acetyl-CoA. Yet, the conversion of acetyl-CoA to fermentative end products was varied. All but one MAG encoded and expressed genes in the pyruvate to succinate pathway. Further, 48/55 MAGs representing all orders except UCB3 transcribed at least one gene for the production of acetate from acetyl-CoA and 32/55 MAGs from all orders except *JAFGEY01* transcribed at least one gene for ethanol production (Figure 3E).

### Anaerobic and Aerobic Respiration Processes

As discussed above, *Anaerolineae* MAGs encoded and expressed genes used in anaerobic fermentation pathways. Other possible anaerobic pathways in salt marshes involve sulfur, and particularly sulfate and/or sulfite, with *Promineifilales* and *Anaerolineales* MAGs recruiting the greatest number of DNA and RNA reads to sulfur cycling genes (Figure 4). Specifically, genes involved in assimilatory sulfate reduction and dissimilatory sulfate reduction had the highest DNA read recruitment, with the reverse order for RNA read recruitment (Figure 4). Out of 55 MAGs, only one MAG encoded a complete dissimilatory sulfate reduction pathway, while a second MAG encoded a dissimilatory pathway for reducing sulfate to sulfite and one MAG was missing the intermediate step in the dissimilatory sulfate reduction pathway. The most common gene encoded and transcribed by MAGs was sulfate adenylyltransferase, present in 53/55 MAGs.

**Figure 4.**
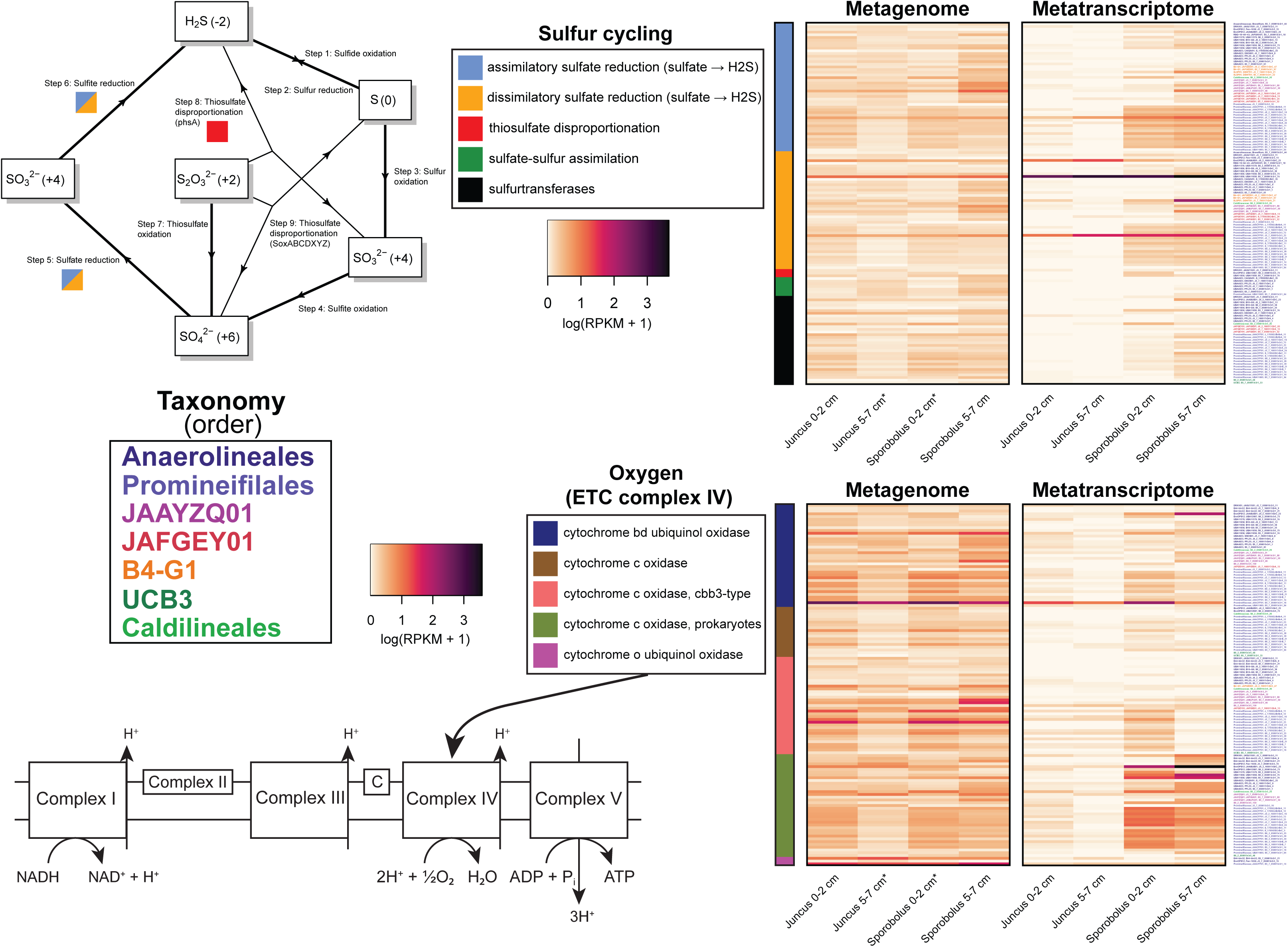
Gene and transcript abundances determined by mapping metagenome and metatranscriptome reads from both plant species (*Juncus* and *Sporobolus*) and depths (0–2 cm and 5–7 cm) to annotated genes and transcripts identified as part of sulfur and oxygen respiratory pathways from the 55 *Anaerolineae* MAGs. RPKM data was log1p-transformed. Genes that can be considered part of more than one pathway have been included in all applicable pathways. An asterisk (*) indicates that RPKM values from two metagenomes have been averaged together prior to transformation.

While the salt marsh rhizosphere microbiome provides a recognized ecosystem service in the reduction of nitrate, no nitrate reduction genes were present in any *Anaerolineae* MAGs recovered in the present study. However, nitrite reduction was encoded in 22/55 MAGs. In fact, the majority of nitrogen cycling genes (21/30 genes) were nitrite reductases. The abundance of nitrogen-cycling genes within the salt marsh *Anaerolineae* MAGs was limited; just 23/55 MAGs were found to encode at least one nitrogen-cycling gene. The greatest number of DNA reads were recruited by the *Anaerolineales* followed by *Promineifilales*. Highest RNA read recruitment was found among the JAFGEY01 and B4-G1, with JAFGEY01 recruiting 1.78-fold more RNA reads than B4-G1.

In addition to the respiratory pathways discussed above, subunits of complex IV, the last step of the electron transport chain (ETC), where oxygen is reduced, were identified in 47/55 MAGs. Of those MAGs, 39 possessed all subunits of at least one complex IV enzyme. Specifically, metagenomic and metatranscriptomic read recruitment to complex IV subunit genes was highest among the *Promineifilales* (152.70 DNA and 63.93 RNA), followed by the *Anaerolineales* (Figure 4). Genes in complex IV encoded and transcribed by the *Anaerolineae* MAGs included those coding for cytochrome bd ubiquinol oxidase, cytochrome o ubiquinol oxidase, and several cytochrome c oxidases, including prokaryote-specific cytochrome c oxidases and a cbb3-type cytochrome c oxidase. In addition, the majority of *Anaerolineae* MAGs appear to encode a suite of complex IV cytochromes; of the 47 *Anaerolineae* MAGs that were found to encode at least one complex IV cytochrome, 41 possessed subunits from at least two different types (Figure 4). Of these genes, genes encoding cytochrome o ubiquinol oxidase recruited the most DNA reads (RPKM of 4.04), followed by cytochrome bd ubiquinol oxidase genes (Figure 4). In contrast, the highest number of metatranscriptomic reads was recruited by prokaryote-specific cytochrome c oxidase genes (RPKM of 2.41) followed by cytochrome bd ubiquinol oxidase genes (Figure 4). All MAGs recruited 0.28-fold more DNA reads and 2.83-fold more RNA reads of complex IV subunit genes from *Sporobolus* samples than *Juncus* samples, and 0.27-fold more DNA and 2.12-fold more RNA reads from *Sporobolus* 5–7 cm samples compared to all other samples. In addition to ETC complex IV, further analysis revealed that 31/55 MAGs encoded all subunits of ETC complex I (NADH:quinone oxidoreductase), 24/55 MAGs encoded all subunits of ETC complex II (prokaryote-specific succinate dehydrogenase), 34/55 MAGs encoded 2 of the 3 subunits of ETC complex III (cytochrome bd ubiquinol oxidase), and 34/55 MAGs encoded at least 7 of the 8 subunits of ETC complex V (F-type ATPase found in prokaryotes and chloroplasts) (data not shown).

### Pathways Encoding Secondary Metabolite Production

Analysis of the 55 *Anaerolineae* MAGs revealed a total of 213 biosynthetic gene clusters (BGCs) were encoded. Of these, 113 BGCs were found on contigs greater than 10 kb in length. When MAGs were grouped by order, the *Promineifilales* were found to have the highest number of total predicted BGCs (90) as well as the highest number of predicted NRPS/PKSs (25), followed by the *Anaerolineales* with 54 predicted BGCs and 20 predicted NRPS/PKSs (Figure 5).When BGCs were grouped by predicted product, 93 encoded terpene synthesis, 66 were predicted to synthesize nonribosomal peptide synthetases (NRPS) or polyketide synthases (PKS), 24 were predicted to synthesize ribosomally synthesized and post-translationally modified peptides (RiPPs), 24 were predicted to contain RiPP recognition elements (RRE), 3 were predicted to synthesize arylpolyene, and 3 were predicted to synthesize indole.

**Figure 5.**
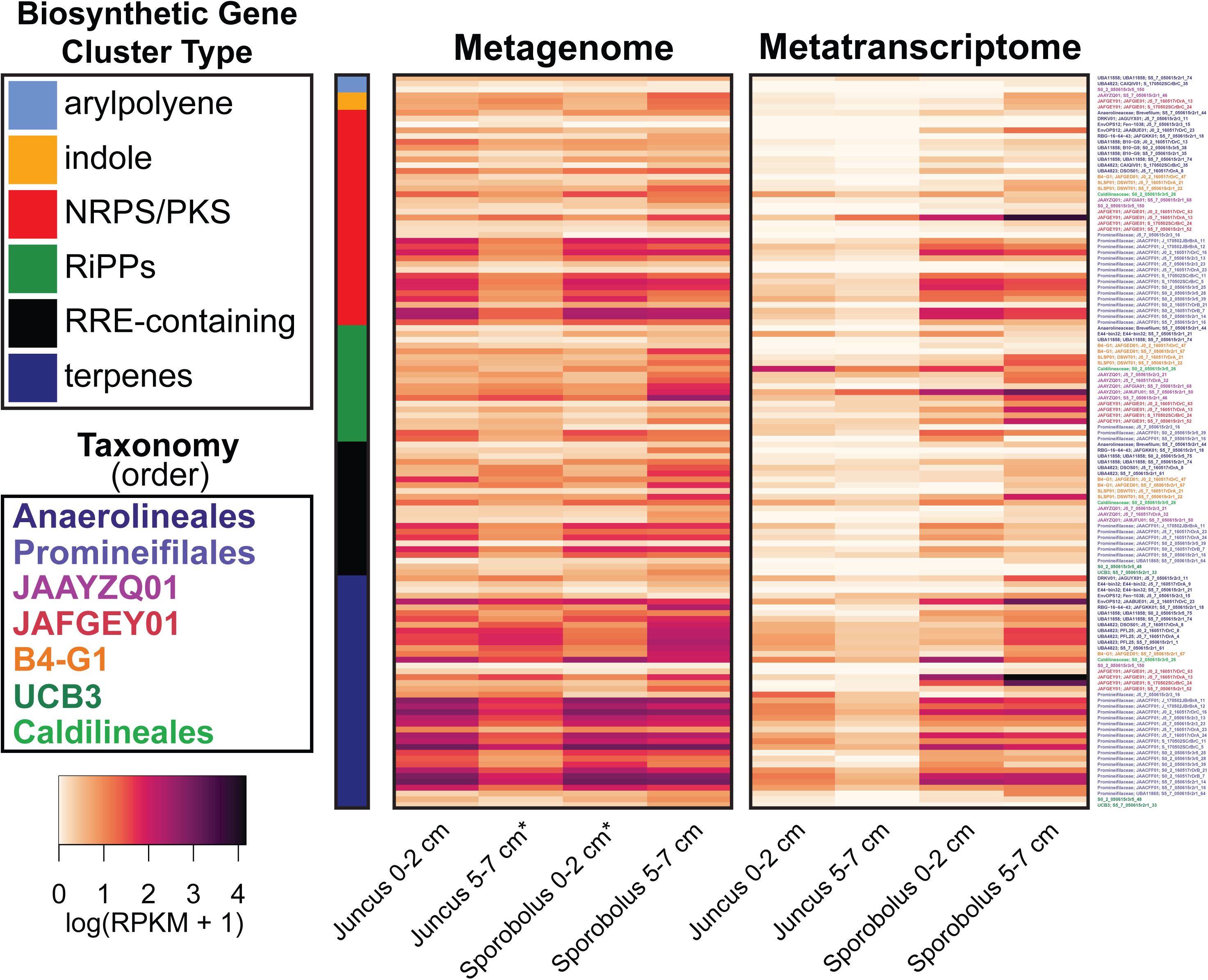
Gene and transcript abundances determined by mapping metagenome and metatranscriptome reads from the 55 *Anaerolineae* MAGs, recruited from both plant species (*Juncus* and *Sporobolus*) and depths (0–2 cm and 5–7 cm), to annotated genes and transcripts identified as biosynthetic gene clusters (BGCs) using Prodigal and antiSMASH. RPKM data was log1p-transformed. BGCs that were identified as belonging to more than one category have been included in both applicable categories. An asterisk (*) indicates that RPKM values from two metagenomes have been averaged together prior to transformation. Some BGC classes have been consolidated into larger groups: cluster type NRPS/PKS contains NRPS, NRPS-like, PKS, hglE-KS, and NRPS-PKS hybrid gene clusters; the cluster type RiPPs contains lanthipeptides, lassopeptides, linaridins, ranthipeptides, RiPP-like, and RiPP-containing hybrid gene clusters.

Mapping of metagenomic and metatranscriptomic reads from identified BGCs to *Anaerolineae* MAGs revealed that the *Promineifilales* recruited the highest number of both metagenomic and metatranscriptomic reads (RPKM of 816.68 DNA and 240.97 RNA) followed by *Anaerolineales* and JAFGEY01 (Figure 5). When recruited reads were summed by BGC type, terpene-synthesizing clusters recruited the greatest number of metagenomic and metatranscriptomic reads (749.76 DNA and 336.96 RNA) followed by NRPS/PKS clusters, which also included NRPS-like, heterocyst glycolipid synthase-like PKS (hglE-KS), and NRPS-PKS hybrid clusters (Figure 5). Grouping BGC read recruitment by grass species revealed that MAGs recruited 0.46-fold more metagenomic reads from *Sporobolus* than from *Juncus* and 2.63-fold more metatranscriptomic reads from *Sporobolus* than from *Juncus*. As with all other pathways analyzed, MAGs recruited the greatest number of metagenomic and metatranscriptomic reads from *Sporobolus* 5–7 cm samples (Figure 5).

## Discussion

Metagenome assembly efforts led to reconstruction of 55 MAGs belonging to seven different orders of *Anaerolineae*. Specifically, we assembled MAGs belonging to the orders *Promineifilales, Anaerolineales,* JAAYZQ01, B4-G1, JAFGEY01*, Caldilineales,* and UCB3 (presented in order of highest to lowest DNA and RNA read recruitment). ANI analysis revealed that in a few cases, MAGs were similar enough to be classified as the same species, although generally MAGs were distantly related (≥59.1% similarity). Metagenomic and metatranscriptomic read recruitment to *Anaerolineae* MAGs showed that *Anaerolineae* were abundant in all samples but were particularly abundant in *Sporobolus* 5–7 cm samples, which agrees with 16S rRNA data presented in Mason et al. (2021) (21). Furthermore, we compared assembled 16S genes from several of our MAGs to 16S gene sequences of *Anaerolineae* presented in Mason et al. (2021) and found that our MAGs were similar to dominant and core members of the microbial community described therein, suggesting that our assembled MAGs captured the abundant *Anaerolineae* as determined with iTag sequencing (21).

*Anaerolineae* were initially described as obligately anaerobic heterotrophic fermenters (2), but more recently diverse metabolic strategies have emerged as a result of omics based analyses (5,6,14). For example, *Anaerolineae* have been reported as prolific degraders of a variety of organic substrates, not only using fermentative pathways, but also utilizing aerobic and anaerobic respiration pathways. Yet, the function of *Anaerolineae* in a salt marsh context is uncharacterized, despite the fact that previous studies have established *Chloroflexota* as a highly abundant member of the salt marsh rhizome microbial community (16–18,20,21).

Sugars make up nearly half of the carbon released by plant roots into the rhizosphere, with hexoses such as glucose dominating over pentoses in soil (64,65). Functional gene analyses of *Anaerolineae* MAGs revealed simple sugar consumption, particularly glucose and galactose, was one of the most abundant and highly expressed carbon degradation pathways in the rhizosphere. In particular, *Promineifilales* and *Anaerolineales* non-CAZyme-encoding genes and transcripts were most abundant. Previous work has demonstrated the capacity of the *Promineifilales* to assimilate glucose for both aerobic respiration and fermentation (1). Our data also showed that salt marsh *Anaerolineae* have the ability to degrade a diverse range of sugars including fructose, fucose, xylose, and mannose, which tend to be found at lower concentrations in soil (65). However, gene expression for the degradation of these sugars was lower than that of glucose and galactose. Our findings regarding salt marsh *Anaerolineae* carbohydrate degradation genes are consistent with previous reports of *Anaerolineae* in other environments, such as *Anaerolineae* found within the holobiont of marine sponges, where *Anaerolineae* were also observed to have the capacity to degrade a diverse assortment of monosaccharides including ribose and xylose (5).

Salt marsh *Anaerolineae* also have the capacity to degrade a variety of polysaccharides with genes encoding CAZymes, and these genes were highly expressed. Genes and transcripts involved in the degradation of starch and sucrose were particularly abundant especially among the *Promineifilales*. Amorphous cellulose consumption was also an abundant pathway. Marsh plants including *Sporobolus* have been previously observed to have increased sucrose concentrations under salinity stress, which may help to explain the high abundance and transcriptional activity of these genes (66). Furthermore, previous studies have measured the polysaccharide content of lignocellulose in *Sporobolus* and *Juncus* and found that *Sporobolus* lignocellulose is 80–93% polysaccharide by weight (including both cellulose and hemicellulose) and *Juncus* lignocellulose is about 75% polysaccharide by weight (67,68). These studies have also demonstrated that the cellulose moiety of lignocellulose is preferentially degraded by microbes compared to the lignin moiety (67,68).

Overall, the ability of the *Anaerolineae* to degrade carbon substrates ranging from monosaccharides such as xylose to more complex carbohydrates and disaccharides such as starch and sucrose is consistent with previous investigations into the metabolism of *Anaerolineae* (2,4,6,14,61) that were sampled from environments other than the salt marsh rhizosphere. Transcription of genes coding for simple sugar degradation was higher than that of genes for complex carbohydrate degradation. This could reflect the abundance of the different carbon substrates. For example, monosaccharides are readily available in the rhizosphere as plant exudates, particularly during the highly productive summer months in which our samples were taken, and are typically consumed by microbes within seconds to minutes (64). In contrast, more recalcitrant carbon sources may remain in the soil for hours to days, or even longer (64). Thus, CAZyme gene abundance and expression could increase slightly in less productive months, though not to the degree that might be observed in a salt marsh with more pronounced seasonality as seen in the northeastern U.S.

Despite access to various monosaccharide and polysaccharide carbon substrates in the salt marsh rhizosphere, the *Anaerolineae* MAGs encoded and expressed genes involved in carbon fixation. Specifically, carbon fixation genes and transcripts were highest among the *Promineifilales*, *Anaerolineales* and JAAYZQ01. Little is known about the JAAYZQ01, though one MAG belonging to this order has been previously assembled from a terrestrial mud volcano environment in association with anaerobic methanotrophic archaea (69). Of the three carbon fixation pathways discussed above, the reductive citrate cycle pathway was the most abundant and highly expressed. This cycle has been observed in anaerobic and microaerophilic members of various phyla including *Chlorobiota* (70). In terms of read recruitment the Wood-Ljungdahl carbon fixation pathway was second only to the reductive citrate cycle. The Wood-Ljungdahl pathway has been observed previously in members of *Chloroflexota* (5,6,14,15). Generally, this pathway is found among acetogenic bacteria which live close to the thermodynamic limit (70). We observed only one of the four key genes required for the 3-hydroxypropionate bi-cycle, which was first described in a member of *Chloroflexota* (71), in only one MAG belonging to *Caldilineales*.

Similar to initial observations regarding *Anaerolineae* as fermenters, salt marsh *Anaerolineae* also demonstrated the capacity for and expression of genes involved in fermentation, particularly among the *Promineifilales* and *Anaerolineales*. Specifically, the fermentative pathways with highest DNA and RNA read recruitment were pyruvate to acetyl-CoA and pyruvate to succinate. Fermentation has been demonstrated within the *Chloroflexota* (6,8,14,15) as well as within the *Anaerolineae* specifically (2–4). Previous work also supports acetate (3,4,8,15,61), lactate (3,8,61), succinate (61), formate (4,61), and ethanol (4,8,61) as fermentative end products produced. In particular, the production of acetate by salt marsh *Anaerolineae* may be favorable given that the MAGs also encode the Wood-Ljungdahl pathway for carbon fixation (70).

Beyond fermentation, other anaerobic processes were annotated in the *Anaerolineae* genomes. Specifically, genes coding for sulfite, sulfate, and nitrite reduction were identified. However, only one genome encoded a complete pathway to reduce sulfate to hydrogen sulfide, and no genomes were found to encode a complete denitrification pathway. Based on the genes that were encoded, it is likely that both of these processes are dissimilatory rather than assimilatory. For example, genomes were found to encode dissimilatory-type sulfite reductase as well as nirS and nirK, which have been used as markers to identify canonical denitrifying bacteria (72). Similarly, a complementary paper that assembled MAGs from the salt marsh rhizosphere, including our metagenomes presented here, reported only 14% encoded complete sulfate reduction (73). It is somewhat surprising that so few genomes from this highly abundant salt marsh clade appear to encode for sulfate reduction or denitrification given the environmental conditions. Salt marsh soils are inundated with water at every high tide, and in this environment oxygen is depleted. Denitrification and sulfate reduction have been demonstrated in the salt marsh rhizosphere with high denitrification rates and porewater hydrogen sulfide concentrations being reported (74,75), including in the salt marsh analyzed herein, where maximum hydrogen sulfide concentration was observed at ∼6000 μmol/L in *Sporobolus*-dominated sediment (21). In contrast, the *Anaerolineae* genomes analyzed here appear to be highly specialized for the oxic rhizosphere environment (see below) where oxygen is much more readily available compared to the surrounding soil, only necessitating the use of sulfate/sulfite or nitrite reduction when the soil is flooded for extended periods.

As discussed above, aerobic respiration by *Anaerolineae* is likely an important form of respiration in the salt marsh rhizosphere, with 47/55 MAGs encoding and actively transcribing ETC complex IV enzymes. While some complex IV enzymes such as cytochrome bd oxidase may be used by bacteria to mitigate hydrogen sulfide toxicity (76), many (37/55) of the *Anaerolineae* genomes also encoded for a near-complete ETC. There are some previous reports suggesting aerobic respiration in *Anaerolineae* (5,14,77), which our findings agree with. However, this was surprising given the reduced nature of the belowground in salt marshes. Further, transcription of prokaryote-specific cytochrome c oxidase (coxBACD) complex IV genes was greater than for genes encoding any other complex IV enzyme. This was unexpected given that, compared to other terminal oxidases, coxBACD is considered to have low affinity for oxygen (78). Conversely, the expression of cbb3-type cytochrome c oxidase, which has a comparatively higher oxygen affinity (78,79), was much lower. This suggests that within the rhizosphere, particularly that of *Sporobolus*, oxygen is abundant enough that there is a limited need to transcribe high-affinity cytochrome c oxidases. In *Sporobolus anglicus*, oxic zones approximately 1.5 mm wide surrounding the plant roots were demonstrated, which arise when excess oxygen is pumped to the subsurface of the soil through the plant’s aerenchyma where it may be released through the rhizodermis to the rhizosphere in a process known as radial oxygen loss (80). While *S. anglicus* has more developed aerenchyma systems than *S. alterniflorus*, the latter may still make oxygen available in the rhizosphere through radial oxygen loss, particularly in stands of tall *S. alterniflorus* (81).

Finally, soil microbes are recognized as the primary source of microbial secondary metabolites. For example, soil-derived *Streptomyces* are major producers of medically relevant secondary metabolites since the 20^th^ century (29,82). Our analysis of *Anaerolineae* MAGs revealed numerous BGCs, thus implicating *Anaerolineae* as a source of novel secondary metabolites. This was particularly true for the orders *Promineifilales* and *Anaerolineales*, with high gene abundance and expression of their BGCs. Nearly half of all identified BGCs were predicted to synthesize terpene compounds, which were the most abundant genes and transcripts for the secondary metabolites that were observed. The terpenoid class of compounds is large and incredibly diverse, with some terpenes being utilized in treatments against diseases such as malaria (83). Until relatively recently, bacteria were not thought to be capable of terpenoid production (83). As a result, terpenoid production by bacteria in general is an emerging but understudied topic. Thus, salt marsh soil should be evaluated as a potential source of novel terpenoid compounds, likely of *Anaerolineae* origin. Many of the BGCs predicted to encode NRPS/PKSs were also identified within the *Anaerolineae* genomes, with abundant NRPS/PKS genes and transcripts. This group of products is of particular interest as some NRPS/PKSs have been identified as medically relevant antibiotics, antifungals, and immunosuppressants (34,84). While other classes within *Chloroflexota* have been previously investigated for their potential as secondary metabolite producers, namely the *Ktedonobacteria* (29), *Anaerolineae* may also represent an untapped source of secondary metabolites with beneficial uses in a soil matrix that is understudied and overlooked.

The uncultured, yet abundant salt marsh *Anaerolineae* analyzed herein encoded and expressed a diversity of carbon degradation and carbon fixation pathways, establishing this group as a primary player in salt marsh carbon cycling. Importantly, the increased read recruitment of carbon degradation genes in *Sporobolus* compared to *Juncus*, as well as the increased read recruitment to ETC complex IV genes, may suggest that soil carbon could be respired more efficiently in the *Sporobolus* rhizosphere than in the *Juncus* rhizosphere by the abundant *Anaerolineae*. Put another way, due to the decreased activity of *Anaerolineae*, which is abundant in association with both plant types, in the *Juncus* rhizosphere, comparatively more carbon is potentially being stored there. From a climate and salt marsh restoration perspective, this may be valuable information. One common method used in salt marsh restoration projects is the replanting of native vegetation (83), which in the southeastern US may include one or both of *Juncus* and *Sporobolus*. These new insights into carbon cycling by the highly abundant *Anaerolineae* in the rhizospheres of each plant may suggest that where possible, *Juncus* may be the optimal choice for replanting due to the potential for higher amounts of carbon sequestration, a clear benefit in the face of anthropogenic climate change. Further, in this salt marsh, greater denitrification rates in this salt marsh were reported in *Juncus* compared to *Sporobolus* (21). Denitrification, or nitrate removal by microbes, is an important ecosystem service, mitigating nitrate input to the marine environment. Collectively, our data and previous reports suggest multiple benefits to replanting with *Juncus*. Regardless of plant type, the intrinsic value of salt marshes has long been recognized in terms of ecosystem services, such as soil carbon storage, providing habitat, flood protection and as discussed above, nutrient filtering, several of which are mediated by the sediment microbiome associated with marsh vegetation. Yet, one overlooked, and potentially important salt marsh function, may be its repository for microbial secondary metabolite production pathways that are encoded in the rhizosphere microbial community. Given, the current global crisis in microbial resistance to anti-microbials, looking to salt marsh soils as a potential source of novel, important secondary metabolites, such as terpenoids, antibiotics, and antifungals, likely of *Anaerolineae* origin, should be considered a priority.

## Acknowledgments

We would like to thank A. Kleinhuizen and L. Linn with assistance in the laboratory. We would also like to thank Philip Hugenholtz and Gene Tyson for the opportunity to analyze metagenomic data at the Australian Centre for Ecogenomics. This project was funded by the National Science Foundation’s Division of Chemical, Bioengineering, Environmental and Transport Systems grants 1643486 (OUM) and 1438092 (BM). Metagenomic sequencing was provided by the Joint Genome Institute through a small-scale community sequencing project grant 503678 (OUM).

## Availability of data and material

Data is available in the appropriate repositories.

